# Robust Generation of Transgenic Mice by Hypotonic Shock Mediated Transgene Delivery in Testicular Germ Cells

**DOI:** 10.1101/049239

**Authors:** Abul Usmani, Nirmalya Ganguli, Subodh K Jain, Nilanjana Ganguli, Rajesh Kumar Sarkar, Mayank Choubey, Mansi Shukla, Hironmoy Sarkar, Subeer S Majumdar

## Abstract

Our ability to decipher gene sequences has increased enormously with the advent of modern sequencing tools but the ability to divulge functions of new genes have not increased correspondingly. This has caused a remarkable delay in functional interpretation of several newly found genes in tissue and age specific manner, limiting the pace of biological research. This is mainly due to lack of advancements in methodological tools for transgenesis. Predominantly practiced method of transgenesis by pronuclear DNA-microinjection is time consuming, tedious and requires highly skilled persons for embryo-manipulation. Testicular electroporation mediated transgenesis requires use of electric current to testis. To this end, we have now developed an innovative technique for making transgenic mice by giving hypotonic shock to male germ cells for the gene delivery. Desired transgene was suspended in hypotonic Tris-HCl solution (pH 7.0) and simply injected in testis. This resulted in internalization of the transgene in dividing germ-cells residing at basal compartment of tubules leading to its integration in native genome of mice. Such males generated transgenic progeny by natural mating. Several transgenic animals can be generated with minimum skill within short span of time by this easily adaptable novel technique.

## Introduction

Transgenesis is an indispensable technology in biomedical research as it allows to manipulate the genome of an organism at will, providing a platform for determining functions of various genes. With the advent of high throughput sequencing technologies, valuable information about several genes and their spatio temporal pattern of expression have been gathered. The technologies have also led to generation of a huge database of information about potential coding and non-coding regions of the genome that control development and maintenance of an organism^1–3^. However, studies of functional genomics are essential to decipher their roles and to understand how their altered expressions are correlated to physiology, development and diseases. Although several methods of germ line gene transfer are available to make transgenic animals for this purpose, they are tedious, time consuming and expensive. For instance, classical method of gene transfer involves microinjection of nucleic acids into fertilized eggs, typically yielding low success rate of about 10-20%^4,5^. Moreover, this technique is beyond reach of common researcher, largely due to technical complexities in adopting skills for micro manipulation of embryos^4^. This has caused a remarkable delay in functional interpretation of several newly found genes in tissue and age specific manner, limiting the pace of biological research. For meaningful interpretation of rapidly generating knowledge about genomics, it is imperative to establish a faster alternative to DNA microinjection mediated transgenesis which is easily adaptable, less invasive and less time consuming.

Alternative to above strenuous technique we had developed a method for generation of transgenic mice by directly injecting the desired gene in the testis followed by in vivo electroporation^6^. Since, this technique requires expertise in survival surgeries of animals, we next developed a two-step non-surgical electroporation procedure to generate transgenic mice^7^.

This technique is less complicated for small animal like mice but due to involvement of electric shock it may not be feasible for large animal transgenesis. Moreover, due to much variation in testis size and scrotal thickness the standardization of voltage parameters remains challenging. This led us to further develop an innovative and simple method to make transgenesis easy.

Treatment with hypotonic Tris-HCl solution result in reduced osmolarity and leads to hypotonic swelling of germ cells which eventually kill them with increased hypotonicity^8^ and this kind of hypotonic-swelling in erythrocyte lead to uptake of surrounding molecules such as nucleosides inside the cell^9^. We sought to exploit this property of the germ cell in testis, and hypothesized that a hypotonic Tris-HCl solution at certain hypotonic concentration might allow the germ cells to internalize the surrounding solutes like DNA in vivo without being killed and the sperm produced from transfected germ cells may carry desired DNA fragment (transgene) which can generate transgenic animal.

Motivated by the above hypothesis, and to circumvent the caveats resident in previous procedures, here we report the development of an easy and simple procedure for *in vivo* gene transfer in male germ cells of the testis for producing transgenic mice. Delivery of the transgene into germ cells was achieved by creating a hypotonic environment surrounding spermatogonia upon testicular injection of linearized transgene suspended in the hypotonic solution of Tris-HCl.

Such a procedure enabling researcher’s to generate their own transgenic animals, instead of outsourcing, would drastically minimize the time required for studies of functional genomics and facilitate research involving humanized transgenic models of diseases.

## Results

### Transfection of nucleotides in testis by Tris-HCl Solution

To test our hypothesis that hypotonicity generated through Tris-HCl solution can help to transfect testicular germ cells in vivo, TRITC labelled dUTP (RED-dUTP) molecules was taken as reporter initially. RED-dUTP suspended in Tris-HCl solution was injected into testis of 30 ±2days old male mice. Earlier it was reported by us that testis of 30 ±2 day old mice is suitable for testicular transgenesis^6^. Initially the concentration of Tris-HCl was maintained at 20mM as reported earlier for treatment to germ cells in vitro^8^. No presence of RED-dUTP was observed in the tubules when isolated from testis. We investigated the possibilities of transfection with higher concentration of Tris-HCl (50mM, 100mM and 150 mM concentrations) with suspended RED-dUTP. We observed the presence of Red-dUTP fluorescence in seminiferous tubulues at 100mM and 150mM concentration, isolated from transfected testis (**Suplimentary Fig. 1a**).

### Transgene transfection in testis by Tris-HCl solution

When it was found that Tris-HCl solution can deliver the nucleotides into the testis, linearized pCX-Egfp plasmid DNA, having EGFP reporter under ubiquitous promoter, was taken for further validation. Variation in parameters were tried to achieve the best transfection in testis in vivo. Injection parameters such as amount of plasmid DNA (ranging from 10 μg −30 μg/testis), volume of injection (ranging from 20 μl – 30 μl), number of injections per testis (1-4) and the concentration of Tris-HCl (ranging from 20 mM to 200 mM) were tested. Based on EGFP expression after 30 days of post transfection in testis, the best transfection in our case was observed with injection conditions of 150 mM Tris-HCl, having 12.5 μg of plasmid DNA in a total volume of 25 μl, and two injection sites per testis of 30±2 days old male FVB mice (**Suplimentary Fig 1b** and **Supplementary Table 1**). In the cross-section of transfected testis, EGFP expression was observed in many tubules. The EGFP expression was also found to be germ cells specific, confirmed by co-localization with VASA, a germ cell marker (Fig. 1a). In transfected tubules there are regions where transfected and non transfected germ cells can be observed (Fig. 1b). We observed no apparent adverse effect of Tris-HCl on transfected testicular tissue architecture (**Supplementary Fig 1c**).

**Fig. 1.**
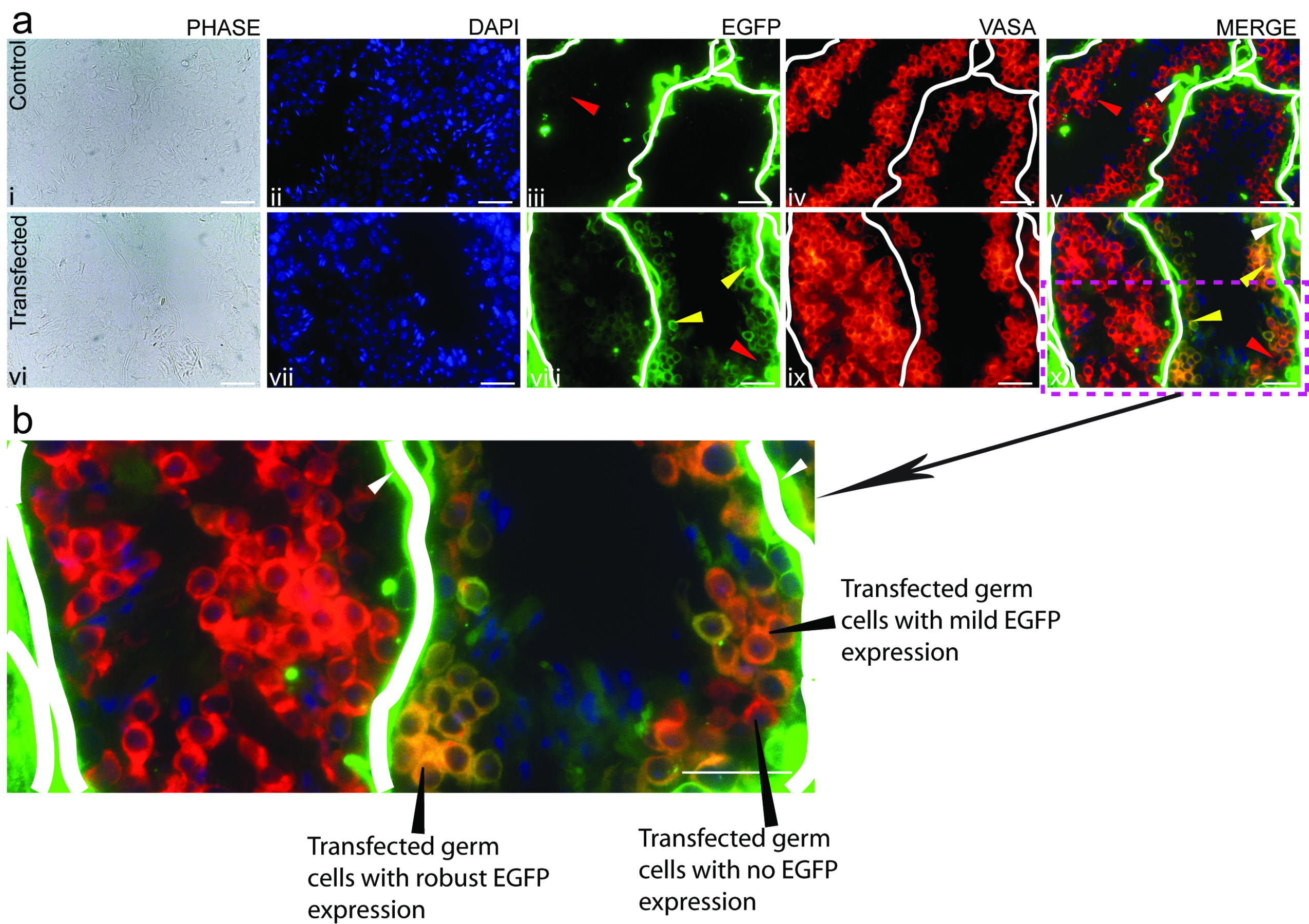
**Transfection of transgene in the testis by hypotonic solution** **a)** EGFP expression in transfected testis transfected with pCX-EGFP suspended in 150mM Tris-HCl. i - v: Tissue Section of non-transfected testis. vi - x: Tissue section of transfected testis. EGFP expression was observed mostly in germ cells (yellow arrow head) of transfected testis. Lack of EGFP expression in non-transfected tubules and germ cells of treated testis (red arrow head). Note: Non-specific signal (white head) in the interstitial space of the testis was observed in both treated and wildtype animal’s tissue section. Scale bar: 50 μm. **b)** Seminiferous tubule showing EGFP in germ cells at higher magnification of section figure - a x. Scale bar: 50 μm.

### Generation of transgenic mice mediated by Tris-HCl testis transfection

Based on our preliminary observations of Tris-HCl mediated transfection of testicular germ cells, and stable expression of transgene even after 30 days of post transfection, we expected that the sperm produce from the transfected germ cells should carry the transgene and could be used for generation of transgenic mice.

Implementing this new method we have generated transgenic mice using *Bucsn2-IRES2-Egfp* construct, which contain *Egfp* reporter gene under Buffalo Beta-casien (Bucsn2) promoter, which expresses specifically in mammary epithelial cells. The linearized construct of *Bucsn2-IRES2-Egfp* was suspended in 150mM Tris-HCl and injected in the both testis of a mice (forefounder) at the age of 30+2 days. Fore-founders were cohabitated with wild type female mice 30 days post transfection to obtain generation one (G1) progeny. The presence of transgenic pups in G1 was detected by PCR (Fig. 2a). Transgenic female mice were housed until adulthood and put for mating to make them lactating. In such transgenic female mice, an intense endogenous EGFP fluorescence was seen in their mammary glands during period of lactation (Fig. 2b). The presence of EGFP was observed in the mammary tissue extract compared to wild type control, and no EGFP expression was found in other tissue types of the same transgenic female mice (Fig. 2c). To evaluate whether the transgene in G1 can be carried forward to the next generation that is G2, transgene positive adult male and female mice from G1 were put for mating. PCR screening showed transgene positive G2 progeny indicating that the transgene was propagated to G2 (Fig. 2a).

**Fig. 2.**
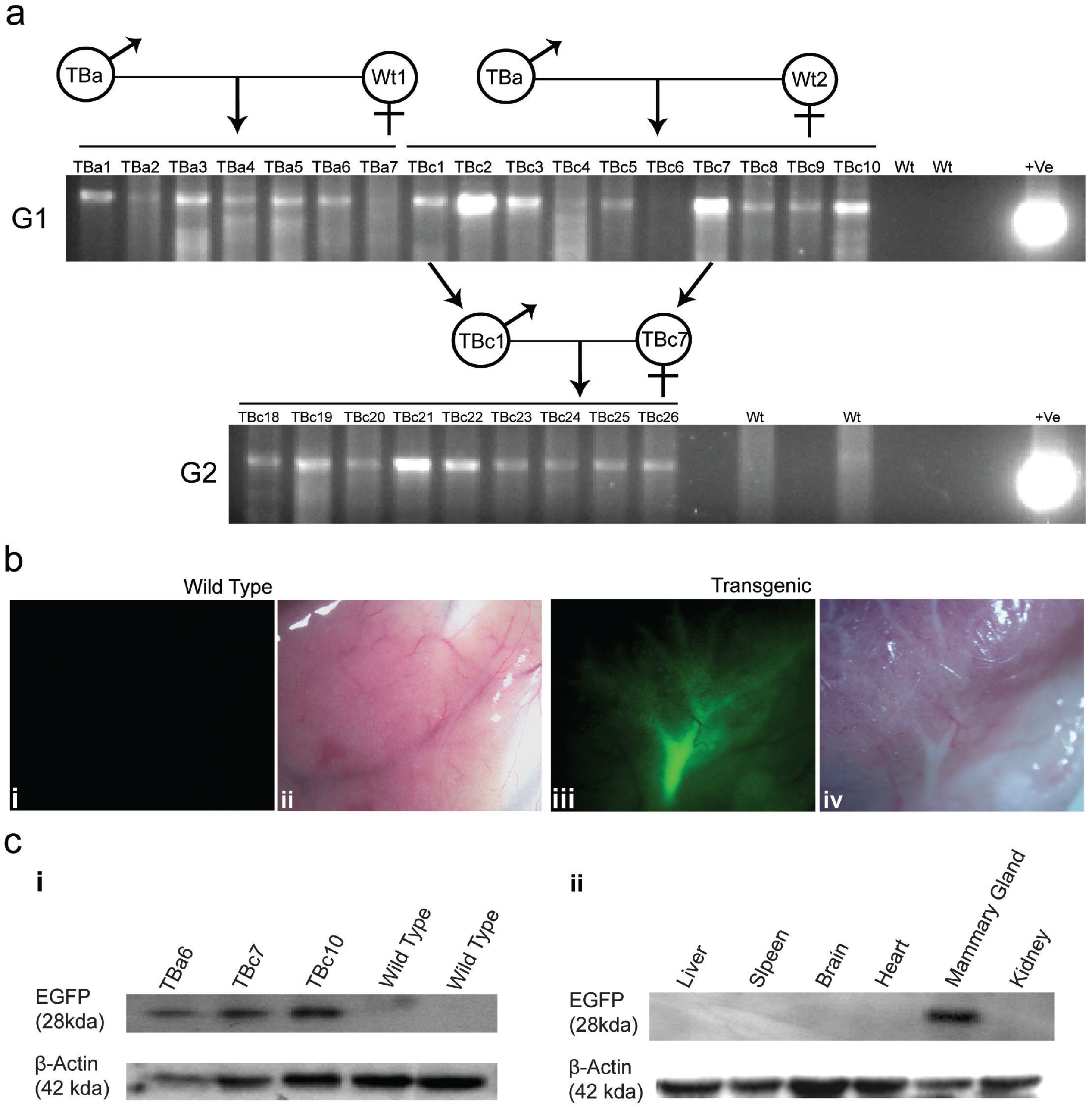
**Generation of transgenic animal with *Bucsn2-IRES2-Egfp* construct** **a)** PCR genotyping of the offspring from generation one (G1) and generation two (G2) to detect transgenic animal. **b)** Mammary gland of lactating (day 7 of lactation) transgenic female mice (TBc 7) carrying *Bucsn2-IRES2-Egfp* transgene, as observed under stereozoom fluorescence microscope. Image (i) show the EGFP fluorescence specifically in mammary gland. Image (iii) show no EGFP fluorescence in gland of wild type mice. Image (ii) and (iv): corresponding phase contrast image. **c)** Expression of GFP by western blot in protein from i) mammary glands of three different transgenic females and two different wild type mice and ii) various tissues (liver, spleen, brain, heart, mammary gland & kidney) of lactating (day 7 of lactation) transgenic female mice (TBc 7) carrying *Bucsn2-IRES2-Egfp* transgene GFP expression (~28 kDa) was observed in transgenic mice whereas there is no GFP signal in wild type mice. GFP expression (~28 kDa) was present only in mammary gland sample and not in other tissue type of the animal. β-actin was used as loading control.

To further validate this newly developed method of transgenesis, we used two more constructs *Amh-IRES2-Egfp* and FetuinA-shRNA for generation of transgenic mice.

*Amh-IRES2-Egfp* construct carried *Egfp* under promoter of anti-mullerian hormone *(AMH)* which is known to be expressed specifically in infant Sertoli cells. By PCR analysis transgene positive animals were detected in G1 (Fig. 3a). Southern blot analysis were performed to determine transgene integration sites in the genome, and it was found that in this transgenic mice line, there were mostly two types of integration of the delivered transgene in the genome. One is integration in tandem repeats which generated ~5.3kb band in southern blot and another is single copy integration for which a band at ~4kb was observed in the same (Fig. 3b). Immuno-histochemical analysis showed Sertoli cell specific expression of EGFP in the testis of 5days old transgenic mice (Fig. 3c) and there was an absence of EGFP expression in the other tissues of the same transgenic mice (**Supplementary Fig. 2a**). As the AMH promoter is infant Sertoli cell specific, no EGFP expression was observed in testis at 42 days (post-pubertal) of age (**Supplementary Fig. 2b**). Western blot analysis with testicular extracts of 5 days old transgenic mice from G1 revealed presence of 28 KDa band corresponding to EGFP (Fig. 3d).

**Fig. 3.**
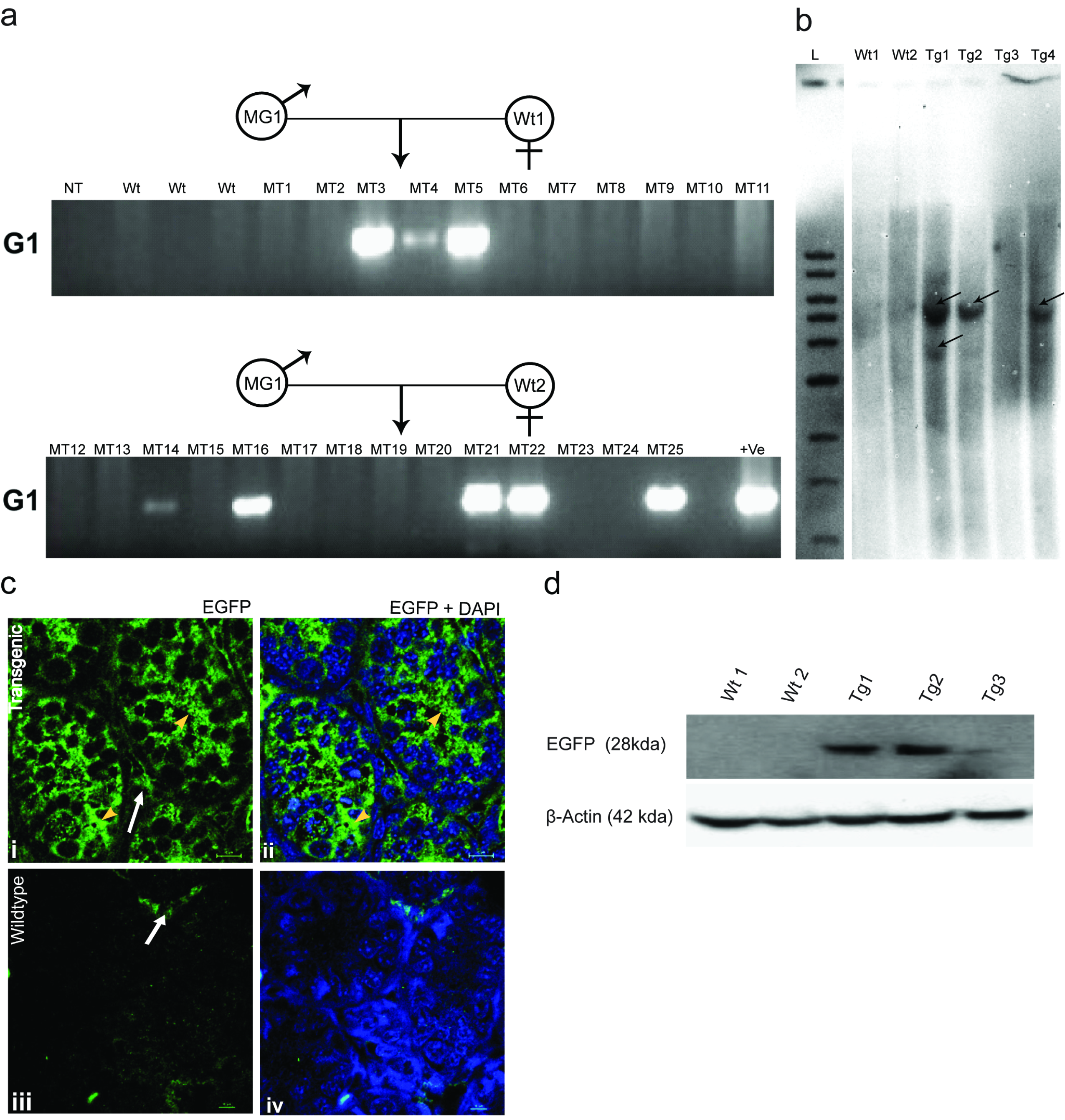
**Validation of *Amh-Ires2-Egfp* transgenic mice**. **a)** PCR genotyping of the offspring using genomic DNA (gDNA) obtained from fore founder MG 1 & MG 2 transfected with *Amh-IRES2-Egfp*. MG 1 & MG 2 were mated with wild type female mice. MG denotes the forefounder animal of *Amh-IRES2-Egfp*. MT denotes transgenic animal of *Amh-IRES2-EgfpE* line. Wt denotes wild type mice. NT denotes no template. +ve denotes plasmid DNA. *G1* = Generation 1, *MT=* Amh-IRES2-Egfp Transgenic. **b)** Southern Blot analysis of genomic DNA isolated from tail biopsy of *Amh-IRES2-Egfp* transgenic mice, showing integration of transgene in multiple sites. wt1 and wt2 denote gDNA isolated from two different wild type animals. Tg1, Tg2, and Tg denotes gDNA isolated from three different transgenic animals. L denotes 1kb DNA ladder (NEB, USA). **c)** GFP expression in Sertoli cells of 5 days old *Amh-IRES2-Egfp* transgenic mice (i, ii) compared with the wild type control. Yellow arrow head shows the EGFP flourescence in the Sertoli cells inside seminiferous tubules. White Arrow marks the nonspecific staining in Leydig Cells. Scale bar: 10μm. **d)**Detection of GFP protein (~28 kDa band) by Western blot analysis from testis of 5 days old three transgenic mice compared with age matched wild-type mice testis. β-actin was used as loading control.

In the construct Fetuin*A*-shRNA, shRNA specific to *Fetuin A* was cloned under ubiquitous promoter U6. The transgenic progeny born in G1-from the fore founders transfected with this construct were screened by slot blot analysis (Fig. 4a) but not with the usual PCR analysis, as in our case the PCR primer did not worked nicely on shRNA may be due to their stable hairpin loop structure. Quantitative Realtime PCR analysis of mRNA extracted from liver tissue revealed that expression levels of Fetuin-A mRNA was drastically reduced in transgenic mice as compared to that in wild type mice (Fig. 4b). Von kossa staining of liver and heart tissue from knockdown mice revealed increased calcification of tissues, unlike wild type tissues (Fig. 4c).

**Fig. 4.**
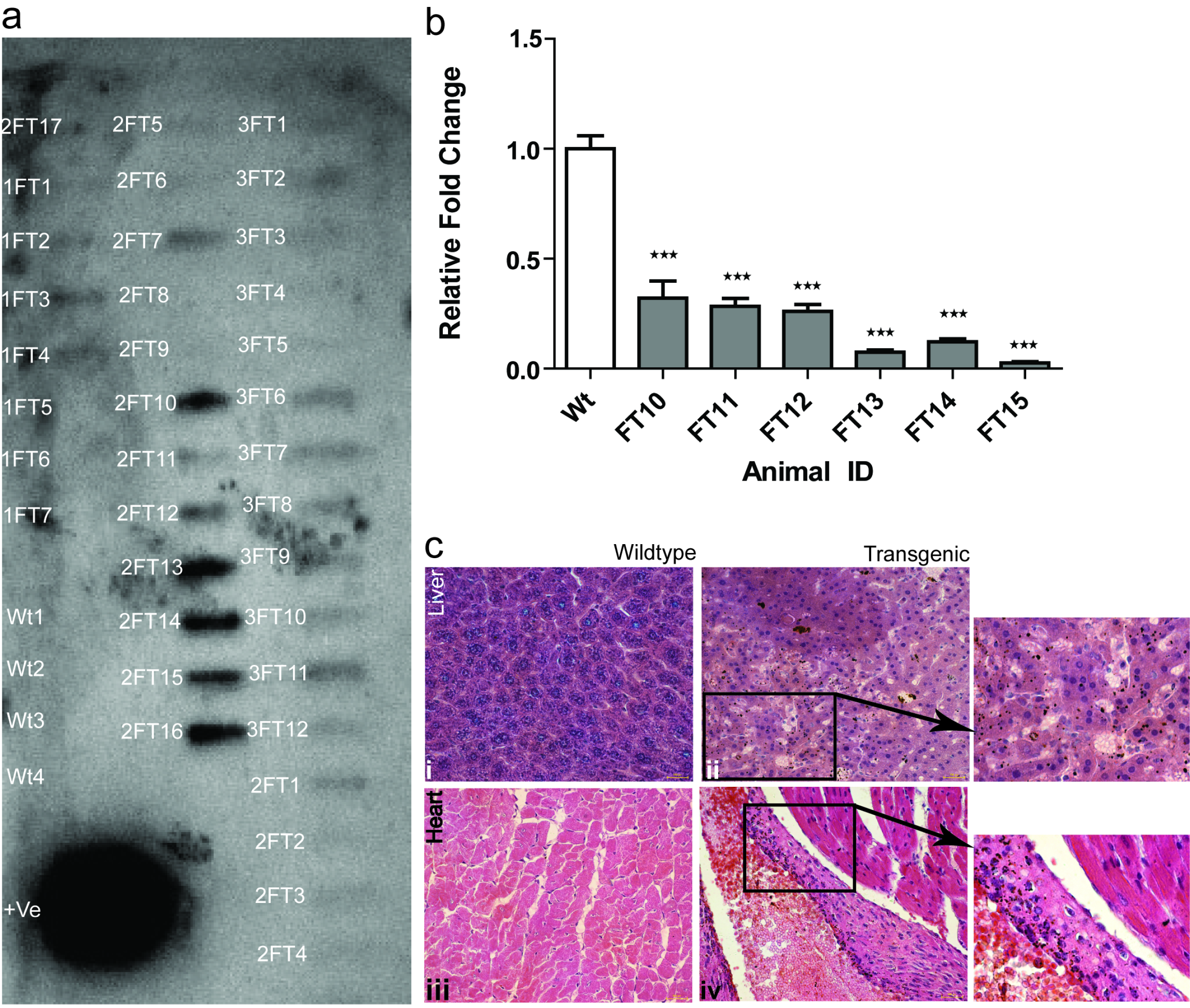
**Evaluation of Fetuin-A knockdown in mice** **a)** Slot blot analysis of the progeny from generation one (G1) obtained from the fore founder males transfected with *Fetuin-A* shRNA. FT – denotes *Fetuin-A* shRNA expressing transgenic mice. Wt – denotes wild type mice. **b)** Relative fold changes in Fetuin-A mRNA expression of transgenic animals relative to wild type animals. wt =wild type mice, F10-F15 represents five different transgenic animals. Each bar generated from n=3 qRTPCR of same sample, represented as mean ± SEM. ^***^ P<0.001. **c)** Von kossa staining for detection of increased calcium deposition (black spots) in Fetuin A knockdown mice. **i)**, **ii)** cross section of liver; **iii)**, **iv)** cross section of heart.

There were 12 out of 17, 8 out of 25, and 16 out of 36 transgene positive animals as judged by genomic analysis of progeny born in G1 generation of *Bucsn2-IRES2-Egfp, Amh-IRES2-Egfp*, and FetuinA-shRNA transgenic mice, respectively. Hence for this newly developed transgenesis technique, the overall efficiency to transmit a transgene in the first generation was found to be 48 % ( **Supplementary Table 2**).

## Discussion

Development of a user-friendly method for rapidly generating transgenic mice without extraordinary laboratory set up is a major unmet need of biomedical researchers. Rapidly generating outflow from modern DNA sequencing technologies has made this need more crucial. To this end, we have developed a technique for making mice which involves simple testicular injection of transgene suspended in hypotonic solution of 150mM Tris-HCl.

Hypotonic solution of 20mM Tris-HCl is usually used to damage and remove the contaminating germ cells from cultures of primary Sertoli cells of testicular origin^8,10^. A treatment with hypotonic solution of 5mM to 20 mM Tris-HCl for 2.5 minutes results into hypotonic-swelling of germ cells^8^. The higher concentration of Tris-HCl (30mM to 80 mM) was inefficient in harming the germ cells^8^. In another study with erythrocytes, hypotonic-swelling led to uptake of surrounding nucleosides, amino acids, and monosaccharides via non-conventional Na^+^– independent pathway^9^. This non-conventional path way suggested to be non-specific for uptake of molecules inside the cells^9^. Here, in this study, we have exploited this cellular response towards hypotonocity and found that hypotonic solution containing Tris-HCl at a concentration of 150mM effectively transfected DNA in testicular germ cells while they being remained healthy. In the process of spermatogenesis, drastic division of testicular germ cells occur with multiple mitotic divisions in the initial phases. In this dividing process genome of the germ cell goes through multiple synthesis phase leading to frequent unwinding of the DNA strands, making it vulnerable for integration and propagation of exogenous DNA fragments, if delivered in to the germ cells. Though there was a chance that other cell types of the testis might also respond to this induced hypotonicity and uptake the delivered transgene, we did not find the EGFP expression in to any other testicular cell type. The hypotonic effect of Tris-HCl does not harm the testis because it gets easily cleared along with the interstitial fluid which has a high turnover rate in the testis^11^.

To determine whether such transfection by hypotonic solution can lead to generation of transgenic animal, we have taken two tissue-specific over expression constructs, namely *Bucsn2-IRES2-Egfp* and *Amh-Ires2-Egfp*, and a knock down construct, U6-shRNA against Fetuin A gene. We successfully generated transgenic progeny by this new method from all the three constructs used in the study. In case of transgenic lactating females carrying *Bucsn2-IRES2-Egfp* transgene, very intense expression of EGFP was observed specifically in the mammary glands. The transgene *(Bucsn2-IRES2-Egfp)* also propagated from generation one (G1) to generation two (G2), further confirming its stable integration in the genome. The results were found to be
similar to our previous report where same construct was used to generate transgenic animals but by electroporation method^12^. In *Amh-Ires2-Egfp* transgenic line, *Egfp* gene was under regulation of murine *Amh* promoter. *Amh* is expressed at very high levels in immature Sertoli cells of the testis from 12.5 day post coitum (dpc) in the mouse^13^ until the onset of testicular puberty^14^ (around 10 days of postnatal age). In transgenic mice carrying *Amh-Ires2-Egfp*, we could successfully detect the pattern of gene integrations in the genome by Southern blot analysis. Age specific expression of EGFP under control of *Amh* promoter in the Sertoli cells of 5 days old mice suggested successful expression of the integrated gene. This was further authenticated by Western blot analysis, which showed the presence EGFP in the testicular extracts from 5 days old transgenic mice but not in the extracts of testis from age matched wild type mice.

We have also evaluated the utility of this method in exerting RNA inhibition *in vivo*. Gene knockdown mice was generated using a construct carrying shRNA against mRNA of Fetuin-A, a protein predominantly produced by the liver. Such mice displayed a significant (p<0.05) decline in the levels of Fetuin-A mRNA in the liver, suggesting successful generation of knockdown mice by this procedure. Fetuin-A is a major inhibitor of calcification in soft tissues, especially that of heart, kidney and lung. Deficiency of Fetuin-A is known to be associated with dystrophic calcification of these tissues^15^. We found similar calcification in hepatic and cardiac tissue of our Fetuin-A knock-down mice generated by this procedure. This observation is in line with previous findings reported in Fetuin A knock-out mice generated using embryonic stem (ES) cells^15^.

The percentage of efficiency for getting a transgene positive animal was found to be about 48 % by this method, which was higher than the conventional method of transgensis by pronuclear DNA injection method 10-20%^4,5^, but lower than the earlier method which used gene electroporation in the testis^7^. Considering the simplicity of this hypotonic solution mediated transgenic method which requires no additional instruments and specific skills, we believe that this efficiency is quite sufficient to generate an adequate number of transgenic founder animals each of which can generate a line for any given biological study. Moreover, this innovative method can be extrapolated in large animals species like non-human primates and bovine where the generation of transgenic animal with the existing techniques are very cumbersome.

In conclusion, we have developed a novel and simple method for making transgenic mice avoiding any harsh treatment to animals. This procedure is fast and can be easily adapted by researchers since it does not require any dedicated laboratory, equipment or specialized expertise to handle embryos. Moreover, it does not involve sacrifice of any animal or use of electric pulses for testicular gene transfer making this technique ethically more acceptable. This easy procedure of *in-vivo* transgenesis by simple injection of a suspension of DNA in the testis provides a remarkable scope to biomedical researchers for generating their own transgenic animal models thereby potentially adding pace to the field of functional genomics.

## Materials and Methods

### Animals

FVB/J strain ofmicewas used for the present study. The mice were housed in a climate controlled environment under standard light (14 hour light, 10 hour dark cycle), temperature (23 ± 1 °C), and humidity (50 ± 5%). Animals were used as per the National Guidelines provided by the Committee for the Purpose of Control and Supervision of the Experiments on Animals (CPCSEA), Govt. Of India. Protocols for the experiments were approved by the Institutional Animal Ethics Committee (IAEC), National Institute of Immunology, New Delhi.

### Plasmids

*pCX-Egfp* plasmid was a kind gift from Dr. Y. Junming (University of Tennesse, Memphis, USA), it contains chicken beta actin promoter along with cytomegalovirus transcription enhancer element (CX) and an *egfpgene*.The plasmid was digested with *Sal I* restriction enzyme to obtain a single fragment of 5.5kb whichwasused for the testicular injection duringstandardization of the procedure(**Supplementary Fig. 3a**).

*Amh-IRES2-Egfp* For this construct,632bp upstream region of the anti mullerian hormone (Amh) gene spanning from −1bp to −632bp from transcription start site of mouse Amh gene was PCR amplified from mouse genome and cloned in *pIRES2-Egfp* vector to generate *Amh-IRES2-Egfp* construct. *Amh-IRES2-Egfp* plasmid DNA was digested with *AseI* and *NheI* restriction enzymes, the fragment of interest (5.3kb) had Amh promoter at 5’ end and *egfp* gene towards the 3’ end (**SupplementaryFig. 3b**).

*Bucsn2-IRES2-Egfp:* This construct expresses *egfp* gene under Buffalo beta casein promoter^12^. *Bucsn2-IRES2-Egfp* plasmid DNA was digested with *PstI and SfoI* restriction enzymes. The fragment of interest (~6.8kb) had BuCSN2 promoter at 5’ end and *egfp* gene towards the 3’ end (**SupplementaryFig. 3c**).

*Fetuin-A-* shRNAconstruct: *Fetuin-A* shRNA bacterial clones targeting mouse *Fetuin-A* gene were procured from Sigma-Aldrich, USA. The shRNA sequence was cloned in pLKO.1-puro vector^16^ (Stewart, 2003) between U6 promoter(RNA polymerase III promoter) and central polypurine tract (cPPT). *Fetuin-A* shRNA plasmid DNA was linearized with *NcoI* restriction enzyme to obtain 7.1 kb fragment, which was used to make transgenic mice (**SupplementaryFig. 3d**).

### Preparation of plasmid DNA

Plasmid DNA was isolated from overnight grown culture of E. coli (dh5α) using plasmid DNA isolation kit (Advanced Micro devices, India) and assessed for quality and quantity of DNA was assessed spectrophotometrically. Samples were checked on 1% agarose gel to check for integrity. Plasmid DNA was digested by appropriate restriction enzymes to take out the functional cassette and were purified by gel extraction kit (Qiagen, USA). Purified DNA was also assessed spectrophotometrically and on agarose gel, before injection in to testis.

### Labelled nucleoside RED-dUTP transfection in to testis

Seminiferous tubules of the 30 days old FVB mice testis were injected with dUTP nucleotide (fluorophore labelled) suspended in different concentration of Tris-HCl hypotonic solution (50 mM, 100 mM, and 150 mM) of pH 7.0. After 2 hours, testis were dissected out and teased to expose the tubules out of the tunica albuginea.Tubules were washed thrice with Phosphate buffered saline (PBS) and observed under u.v. for the presence of fluorescence inside the seminiferous tubules.

### Hypotonic solution mediated *in vivo* gene transfer in testicular germ cells

Male mice (30±2 days post birth) were anesthetized using intra-peritonial injection (±120 μl) of amixtureof ketamine (45mg/kg) and xylazine (8mg/kg). Hair from scrotal area were trimmed, followed by disinfection with Betadiene. The area was rinsed after 2 minutes with 70% alcohol, leaving the area clean and moist. Gently, testis were pushed down from abdominal cavity with the help of thumb and index finger. Plasmid DNA suspended in Tris-HCl solution (concentrations ranging from 20mM to 200mM during standardization) of pH 7.0, along with 0. 04% Trypan blue and was injected slowly in to the descended testis using 26 guage10μl volume Hamilton syringe (701N; Hamilton Bonaduz AG, Switzerland). For standardization, a range of 20-30μl of hypotonic solution containing various concentration of linearized plasmid DNA (0.5-1.5μg/μl) was delivered into single testis. Variation in the concentration of Tris-HCl solution, concentration of plasmid DNA and number of injection sites (1-4) were done at the time of standardization of this technique (**Supplementary Fig. 4**).

### Generation and screening of transgenic lines

Two constructs for over-expression and one construct for downregulation by shRNAwere used for development and validation of this new procedure.The injected mice (fore-founders) were put for natural mating with wild type adult females after 30-35 days post-injection (mice age 60-65 days) and genomic DNA of progeny were analyzed for the presence of transgene. For this purpose, tail biopsies were obtained at 21 days of age and genomic DNA of mice was extracted from respective tissue. Presence of transgene in the genomic DNA was determined by PCR using transgene-specific primers (**Supplementary Table 3**). PCR was performed using the standard protocol (Sambrook and Russell, 2001). The PCR products were analyzed by TAE agarose gel electrophoresis. To rule out the possibility of false positives in the PCR, negative controls such as a reaction with the gDNA of the wild-type (FVB/J) mice, were performed. For positive control, a reaction was performed using 20ng of pBuCSN2-IRES2-EGFP plasmid DNA.

Genomic DNA of pups were also analyzed for gene integration by Slot blot analysis for transgene constructs bearing shRNA. In brief, probe identifying the transgene fragment was generated by αP^32^dCTP using High Prime DNA labeling kit (Roche Diagnostic GmbH, Mannheim, Germany). Denatured gDNA (1 μg) was blotted on Hybond N+ (Amersham Pharmacia Biotech, England) membrane with slot blot apparatus (Cleaver Scientific, Warwickshire, UK) under vacuum. The membranes were pre-hybridized for 4hours, followed by hybridization with respective transgene specific probes for 10-12 hours. The hybridized blot was exposed to Kodak BioMax MR Film (Kodak, Rochestar, New York, USA) for detection of signals by autoradiography.

### Southern blot analysis

Southern blot analysis was performed following standard procedure^17^. Ten micrograms of genomic DNA obtained from transgenic progenyof G1 generation was digested with BamHI restriction endonuclease. Digested product was resolved on 1% agarose gel, and transferred to Hybond N+ (GE Healthcare, England). An *Egfp* probe fragment of ~600bpwas labelled with αP^32^dCTP using High Prime DNA labeling kit (Roche Diagnostic GmbH, Mannheim, Germany) and was used to detect the transgene integrations. The membrane was hybridized for 20 hours and was exposed to Kodak BioMax MS film (Kodak, Rochestar, New York, USA) for detection of signals by autoradiography.

### Histology and immuno-histochemistry

For histology, tissues were dissected, fixed in formalin and processed for paraffin embedding. Sections were stained by hematoxylin and eosin or with Vonkossa stain. We determined the expression of EGFP in the seminiferous tubule of *pCX-Egfp* and *Amh-IRES2-Egfp* transgenic mice by immuno-histochemistry. Testis sections of 5μm were subjected to immunostaining with mouse anti-GFP (Clontech, Mountain View, CA, USA) as primary antibody at a dilution of 1:200 and then anti mouseIgG tagged to alexafluor 488 (Molecular probes, Eugene, OR, USA) was used at a dilution of 1:250, as secondary antibody. The fluorescence was detected by Nikon Eclipse Ti inverted fluorescence microscope (Nikon Corporation, Chiyoda-ku,Tokyo, Japan). The images were captured using Nikon-digital sight DS-Ri1 camera.

### Western blot analysis

Tissue lysates from testis of *Amh-IRES2-Egfptransgenic* mice; mammary gland, liver, spleen, brain, heart and kidney ofBucsn2-IRES2-Egfp transgenic mice were used to determine presence of EGFP.30μg of protein sample was resolved on 12% SDS-PAGE and transferred to PVDF membranes (MDI, India). The membranes were first incubated with primary antibody (mouse anti-GFP) at 1:1000 dilutions followed by anti-mouse secondary antibody conjugated with horseradish peroxidase at 1:5000 dilutions. The protein bands were detected using enhanced chemiluminescence method (ECL, Amersham Biosciences, UK).

### RNA isolation and Real-Time PCR

RNA was isolated from liver tissue of *Fetuin-A* knockdown transgenic mice and wild type mice using TRIzol (Sigma Chemical Co., USA). Real time PCR was performed using primer specific for Fetuin-A gene (**5**’TCACAGATCCAGCCAAATGC**3**’ as forward primer and **5**’GGAATAACTTGCAGGCCACT**3**’ as reverse Primer). RNA (1μg) was treated with DNase I (1 unit; Fermentas, Pittsburgh, PA, USA) for 30 minutes at 37°C. Reaction was terminated by adding 1μl of 25 mM EDTA and incubating at 65°C for 10 minutes. DNaseI treated RNA was reverse transcribed using Reverse Transcription System (Eurogentec, Seraing, Belgium) with MuMLV reverse transcriptase enzyme and oligo-dT (15mer) for the single-strand cDNA synthesis. Real time PCR amplifications were performed in the Realplex (Eppendorf, Hamburg, Germany) in a total volume of 10μl, which included 1μl of cDNA, 5μl of Power SYBR Green Master Mix (Applied Biosystems, CA, USA)and 0.5μl of each primer. Expression of GAPDH [using**5**’AGAACATCATCCCTGCATCC**3**’ as forward primer and **5**’CACATTGGGGGTAGGAACAC**3**’ as reverse Primer] was analyzed for use as an endogenous housekeeping gene control. Relative fold change of Fetuin-A mRNA in transgenic animal with respect to wild type mice was calculated by 2^ΔΔct^ method^18^.

## Author Contribution

The first and second authors contributed equally to this work. The experiments were conceived and designed by SSM and AU. Experiments were performed by AU, NG1 (Nirmalya Ganguli), NG2 (Nilanjana Ganguli), RS, MC, HS, MS. The data presented in the manuscript were analysed by all authors. The manuscript was written by AU, NG1 and SSM.

## Acknowledgement

We are grateful to the Director of National Institute of Immunology, for her support. We acknowledge the financial support provided by Department of Biotechnology, Govt. of India, under grants BT/PR11313/AAQ/01/376/2008 and BT/HRD/35/01/01/2010. We are grateful Dr. Y. Junming for providing pCX-EGFP plasmid. We greatly acknowledge the technical support from Birendra N. Roy, Dharamveer and Ram Singh. We acknowledge the help from staff of Small Animal Facility of National Institute of Immunology.

